# Population and species diversity at mouse centromere satellites

**DOI:** 10.1101/2020.10.20.345884

**Authors:** Uma P. Arora, Caleigh Charlebois, Raman Akinyanju Lawal, Beth L. Dumont

## Abstract

Centromeres are satellite-rich chromatin domains that are essential for chromosome segregation. Centromere satellites evolve rapidly between species but little is known about population-level diversity across these loci. We developed a *k*-mer based method to quantify centromere copy number and sequence variation from whole genome sequencing data. We applied this method to diverse inbred and wild house mouse (genus *Mus*) genomes and uncover pronounced variation in centromere architecture between strains and populations. We show that patterns of centromere diversity do not mirror the known ancestry of inbred strains, revealing a remarkably rapid rate of centromere sequence evolution. We document increased satellite homogeneity and copy number in inbred compared to wild mice, suggesting that inbreeding remodels mouse centromere architecture. Our results highlight the power of *k*-mer based approaches for probing variation across repetitive regions and provide the first in-depth, phylogenetic portrait of centromere variation across *Mus musculus*.

## Introduction

Centromeres are chromatin domains that are essential for chromosome segregation and the maintenance of genome stability (Bakhoum, Thompson, Manning, & Compton, 2009; Fukagawa & Earnshaw, 2014; McKinley & Cheeseman, 2016; Schalch & Steiner, 2017). Centromeres serve as focal points for the assembly of the kinetochore complex, which provides the protein interface linking chromosomes to microtubules during mitosis and meiosis (Bakhoum et al., 2009; Fukagawa & Earnshaw, 2014; McKinley & Cheeseman, 2016; Schalch & Steiner, 2017). Mutations that abolish or reduce centromere function can impair kinetochore assembly and lead to spontaneous chromosome loss, cell cycle arrest, or chromosome mis-segregation(Holland & Cleveland, 2009). Thus, the loss of centromere integrity can have adverse consequences for genome stability, and represents an important mechanism leading to both cancer and infertility (Aldrup-MacDonald, Kuo, Sullivan, Chew, & Sullivan, 2016; Barra & Fachinetti, 2018; Hudson et al., 1998; Régnier et al., 2005; Zhang et al., 2016).

In most vertebrate species, centromeric DNA is comprised of tandem arrays of one or more satellite repeat units (Malik & Henikoff, 2009; Rocchi, Archidiacono, Schempp, Capozzi, & Stanyon, 2012; Ventura et al., 2007). As a consequence of this satellite-rich architecture, centromeres are predisposed to high rates of structural mutation via replication slippage, unequal exchange, and transposition (Barra & Fachinetti, 2018). These processes actively contribute to size and sequence variability between species (Alexandrov, Kazakov, Tumeneva, Shepelev, & Yurov, 2001; Alkan et al., 2011; Cacheux, Ponger, Gerbault-Seureau, Richard, & Escudé, 2016; Musich, Brown, & Maio, 1980). For example, in mammals, centromere repeat sizes range from 6bp in the Chinese hamster (*Cricetulus griseus*) to 1419 bp in cattle (*Bos taurus taurus*), with GC content ranging from 28−74% (Melters et al., 2013). The remarkable size and sequence variability of centromeres, combined with their critical and highly conserved cellular roles in chromosome segregation and genome stability, impose an enduring biological paradox.

Due to their inherent repeat-rich nature, centromeres persist as gaps on most reference genome assemblies. To date, only a handful of mammalian centromeres (human chromosomes 8, X, and Y) have been fully sequenced and assembled (Jain et al., 2018; Longsdon et al., 2020; Miga et al., 2020). The near absence of high-quality reference sequences and the challenge of uniquely anchoring short reads within repeat-rich regions pose significant barriers to the discovery and analysis of genetic variation across these functionally critical regions. Consequently, the scope of centromere structural and sequence diversity within and between populations remains largely unknown.

Defining levels of centromere diversity represents a crucial first step towards understanding the potential phenotypic consequences of variation at these loci. Prior studies in humans have identified centromere variants that associate with differences in the stability of kinetochore protein binding, which can, in turn, influence the fidelity of chromosome segregation(Aldrup-MacDonald et al., 2016). Studies in mice and monkeyflowers (*Mimulus*) have shown that differences in centromere size can lead to biased, non-Mendelian chromosome transmission in heterozygotes, a phenomenon known as centromere drive (Chmátal et al., 2014; Fishman & Kelly, 2015; Iwata-Otsubo et al., 2017). However, owing to an incomplete catalog of centromere diversity and the omission of variants in these regions from GWAS and linkage studies (Aldrup-MacDonald et al., 2016; Langley, Miga, Karpen, & Langley, 2019), the contribution of centromere variation to phenotypic variation, including disease, has yet to be fully realized.

House mice (genus *Mus*) provide an ideal system for ascertaining population level centromere satellite diversity and evaluating its functional consequences for several reasons. First, prior investigations have identified the focal *Mus musculus* centromere satellite repeat sequences, and defined core features of house mouse centromere architecture (Kalitsis, Griffiths, & Choo, 2006; Kipling, Wilson, Mitchell, Taylor, & Cooke, 1994; Narayanswami et al., 1992; Wong & Rattner, 1988). Specifically, the *Mus musculus* centromere is composed of two primary satellite domains. The minor satellite domain is a tandem array of a 120-bp sequence that cumulatively extends over ~1 Mb of sequence per chromosome. This satellite array delimits the region where the centromere-specific histone variant CENP-A is bound and defines the core centromere (McKinley & Cheeseman, 2016). The minor satellite region is flanked by a 234-bp major satellite repeat array that extends over ~2 Mb of sequence per chromosome. The major satellite region forms the pericentromeric heterochromatin, which is important for sister chromatid cohesion during cell division (McKinley & Cheeseman, 2016; Peters et al., 2001). Second, mouse centromeric satellite arrays are reported to be homogenous both within and between chromosomes (Kalitsis et al., 2006; Wong & Rattner, 1988), a feature that simplifies the task of quantifying their variation in genomes. This contrasts with the architecture of human centromeres, which are composed of distinct repeat arrays that form higher order repeats that vary between chromosomes (Alexandrov et al., 2001; Musich et al., 1980; Wong & Rattner, 1988). Third, whole-genome sequences from diverse wild and inbred *Mus musculus*, as well as more divergent *Mus* taxa are publicly available (Adams, Doran, Lilue, & Keane, 2015; Harr et al., 2016; Thybert et al., 2018). These resources enable surveys of centromere diversity along several dimensions, including among inbred strains, within natural populations, between subspecies, and between species. Finally, as the premiere mammalian biomedical model system, house mice are equipped with experimental tools and detailed phenotype catalogs that can be leveraged to test the functional consequences of centromere variation.

Here, we harness these strengths of the *Mus musculus* model system to carry out the first sequence-based analysis of centromere diversity and evolution in mice. We couple *k*-mer based bioinformatic methods with experimental approaches to uncover remarkable variation in the size and sequence composition of centromeres across a panel of diverse inbred strains and wild-caught house mice. Overall, our study yields a portrait of centromere satellite diversity across a group of closely related mammals and lays the groundwork for future functional studies on the consequences of natural genetic variation at these essential chromatin domains.

## Materials and Methods

### Whole Genome Sequencing Data

Illumina whole genome sequences from 100 house mouse (*Mus*) genomes were obtained in binary alignment map (bam) and fastq formats from public repositories (Supplementary Table 1). These samples include 33 inbred house mouse strains of predominantly *Mus musculus domesticus* ancestry(Adams et al., 2015), 27 wild *M. m. domesticus* mice from four populations, 22 wild *M. m. musculus* from three populations, ten wild *M. m. castaneus* from India, eight wild *M. spretus* from Spain (Harr et al., 2016), a wild-derived inbred strain of *Mus caroli* (CAROLI/EiJ), and a wild-derived inbred strain of *Mus pahari* (PAHARI/EiJ)(Thybert et al., 2018). *Mus caroli* and *Mus pahari* sequence reads were mapped to the *Mus musculus* reference (mm10) using bwa mem version 0.7.9 (Li & Durbin, 2010). Optical duplicates were removed using the *rmdups* command in samtools version 1.8 (Li et al., 2009).

**Supplementary Table 1: House mouse (*Mus*) whole genome sequence samples.** Numbers represented in the data source column correspond to the following data sources: (1) The Mouse Genomes Project Release 1502/REL-1502 (Adams et al., 2015); (2) (Thybert et al., 2018); (3) Wild Mouse Genomes Project (Harr et al., 2016).

*We excluded libraries from these Sanger inbred strains that generated reads with length <75bp.

### *k*-mer Frequencies and Normalization

We computed the observed frequency of all *k*-mers in each mouse genome on a per-library basis. Briefly, each sequenced read in a sample’s fastq file was decomposed into its constituent nucleotide words of length *k*, or *k*-mers using a custom Python script (KmerComposition.py). We selected two lengths of *k*: *k* = {15, 31}. These *k* values were selected to balance computational speed (*k*=15) and provide high sequence specificity (*k*=31). Each analyzed genome captured 440-965 million unique 15-mers and 1.1 - 14.5 billion unique 31-mers.

The efficiency of PCR amplification is not uniform with respect to GC-content, and this can lead to biases in the nucleotide composition of sequencing libraries (Benjamini & Speed, 2012). If uncorrected, such biases could cause false inference of differences in *k*-mer abundance between independent libraries and samples. We implemented a GC-correction to rescale raw *k*-mer counts by the extent of the observed GC-bias in each library. Briefly, we randomly selected a set of ~100,000 *k*-mers that occur uniquely in the mouse reference genome (mm10). For each sample, we modelled the observed counts of these unique *k*-mers as a function of their GC-content using LOESS regression, with the span parameter set to 0.4. The LOESS regression produced a predicted *k*-mer count for each GC-content bin; these values correspond to the magnitude and direction of the empirical GC-bias in the sequencing library and represent the expected “amplification” of a *k*-mer based on its GC-content. Finally, observed *k*-mer frequencies were normalized by the LOESS predicted count for the corresponding GC-content bin:

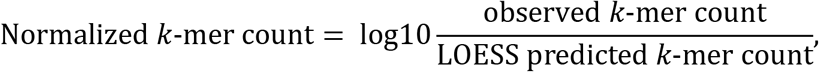

Normalized values were used for comparisons across libraries and samples.

We used reads derived from multiple independent sequencing libraries from a single inbred strain to confirm that our strategy was robust to potential artifacts introduced during library preparation. After GC-correction, we observe excellent concordance in centromere *k*-mer frequencies among replicate libraries for a given strain (Pearson correlation 0.990 < R^2^ < 0.999; Supplementary Figure 1). As expected, concordance between strains was generally weaker (Pearson correlation 0.41 < R^2^ < 0.998; Supplementary Figure 1).

**Supplementary Figure 1:**
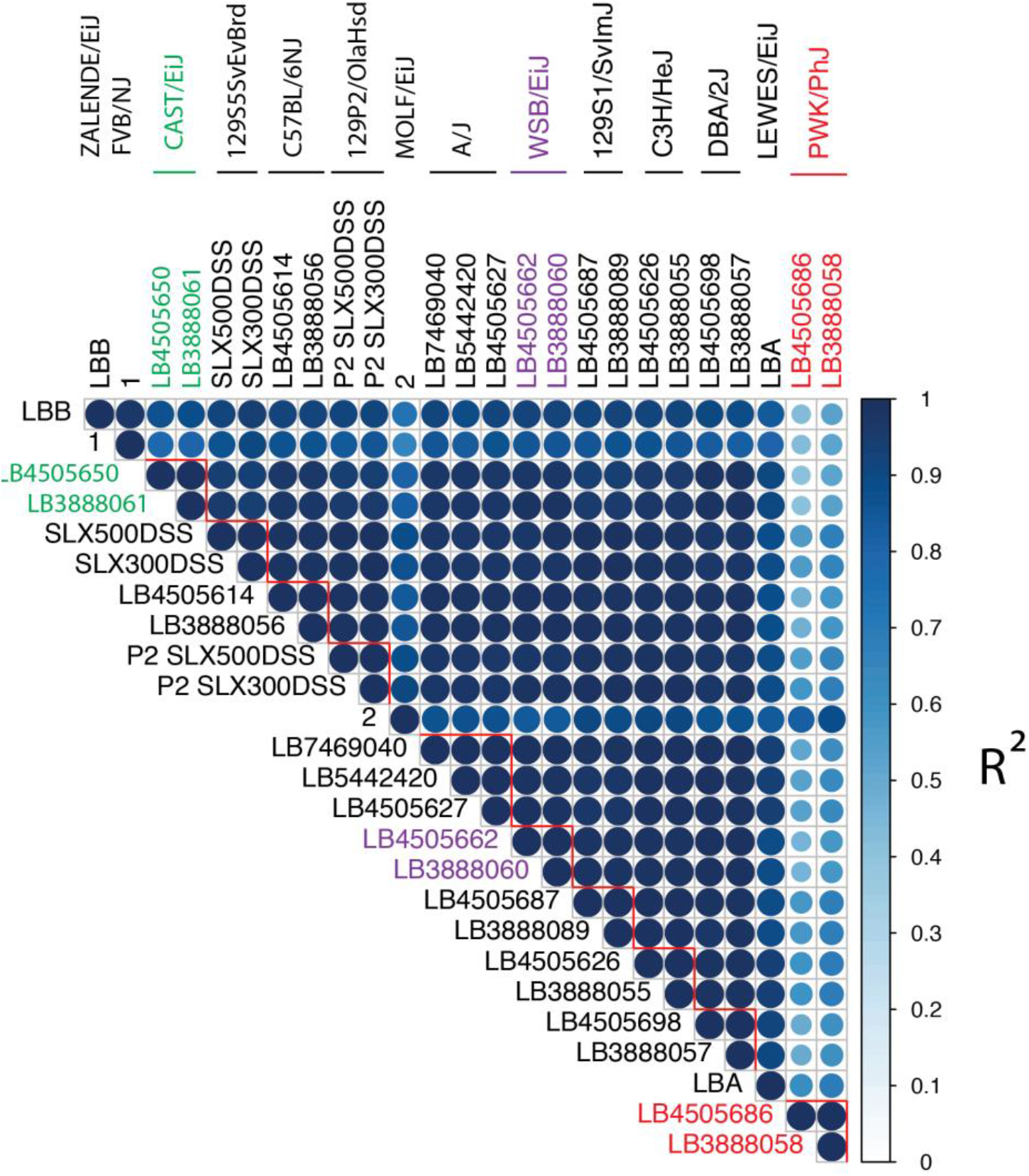
Concordance of GC-corrected *k*-mer counts among strains and replicate libraries within a strain. Heatmap of pairwise Pearson correlations between GC-corrected consensus centromere 31-mer frequencies from replicate sequencing libraries across inbred *Mus musculus* strains. Both color intensity and circle size correspond to the magnitude of the R^2^ correlation coefficient. Red lines delimit replicate libraries for single inbred strains.

### Identification of highly variable *k*-mers across house mouse

To identify *k*-mers that differ in abundance across genomes, we selected a representative subset of *n*=54 diverse *Mus* samples (see Supplementary Table 1) and computed the variance in observed 15-mer frequencies across their genomes:

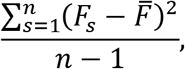

where *F* is the absolute 15-mer frequency standardized by the read depth of strain *s* and 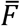 is the average normalized frequency of the 15-mer across the selected 54 strains. The 1000 15-mers with the largest variance were plotted as a heatmap using the R package *pheatmap*. Clusters of closely related 15-mers differing by a single nucleotide offset were manually assembled into longer sequences.

### Quantifying centromere satellite abundance

We used a reference-informed approach to quantify the relative copy number of centromeric satellites in each mouse genome. We first decomposed the minor and major satellite consensus sequences into their constituent *k*-mers (Wong & Rattner, 1988). We then queried the GC-corrected frequency of these centromere *k*-mers in each analyzed library and compared the distribution of these centromere *k*-mer frequencies across libraries and samples.

The relative copy number of a centromere satellite consensus sequence in a given mouse genome was estimated from the median frequency of all constituent *k*-mers. For example, if the median log10 GC-corrected count for *k*-mers present in the major satellite in a given genome was 5, we estimated 10^5^ copies of the major satellite in that genome. This quantity is highly correlated with the overall percentage of sequenced reads that map to the minor and major consensus sequences (Pearson correlation; minor: R^2^ = 0.73, P=2.64×10^−5^; major: R^2^ = 0.85, P=1.65×10^−7^; Supplementary Figure 2), suggesting that it provides a faithful readout of centromere satellite copy number.

**Supplementary Figure 2:**
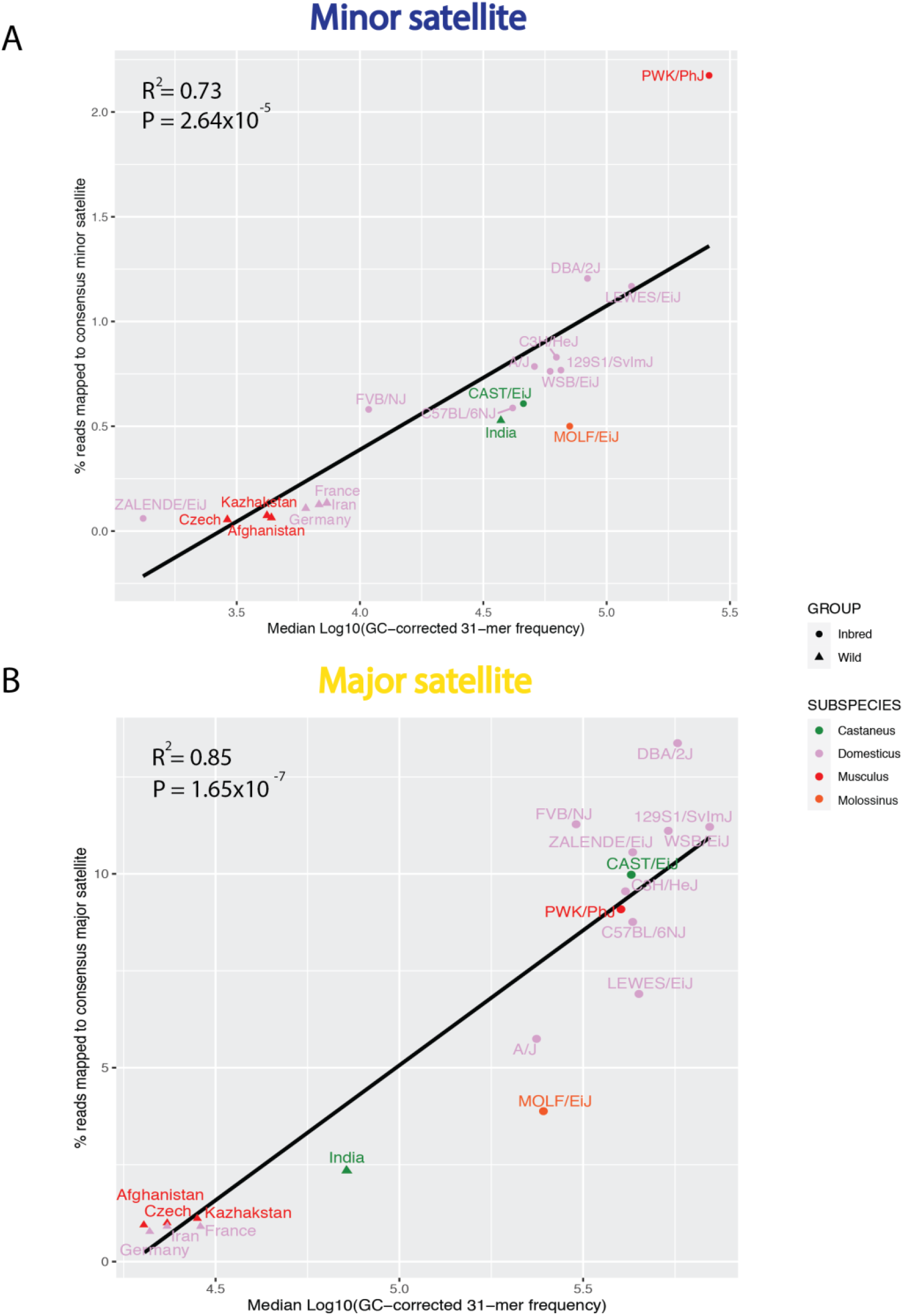
Centromere consensus 31-mer estimates of relative copy number strongly correlate with the percentage of reads mapping to the centromere consensus. Correlation plots for median GC-corrected centromere consensus 31-mers frequency and the percentage of reads mapping to the centromere consensus for the (A) minor and (B) major satellite. Subspecies are represented by color. Inbred and wild-caught mice are distinguished by shape.

We observe little variation in centromere satellite copy number among wild-caught mice sampled from a single population and replicate sequencing libraries from a single inbred strain (Supplementary Figure 3). We therefore combine all individuals from a given population to produce a single population-level copy number estimate. Similarly, for inbred strains with multiple sequencing libraries, GC-corrected *k*-mer counts were aggregated across libraries.

**Supplementary Figure 3:**
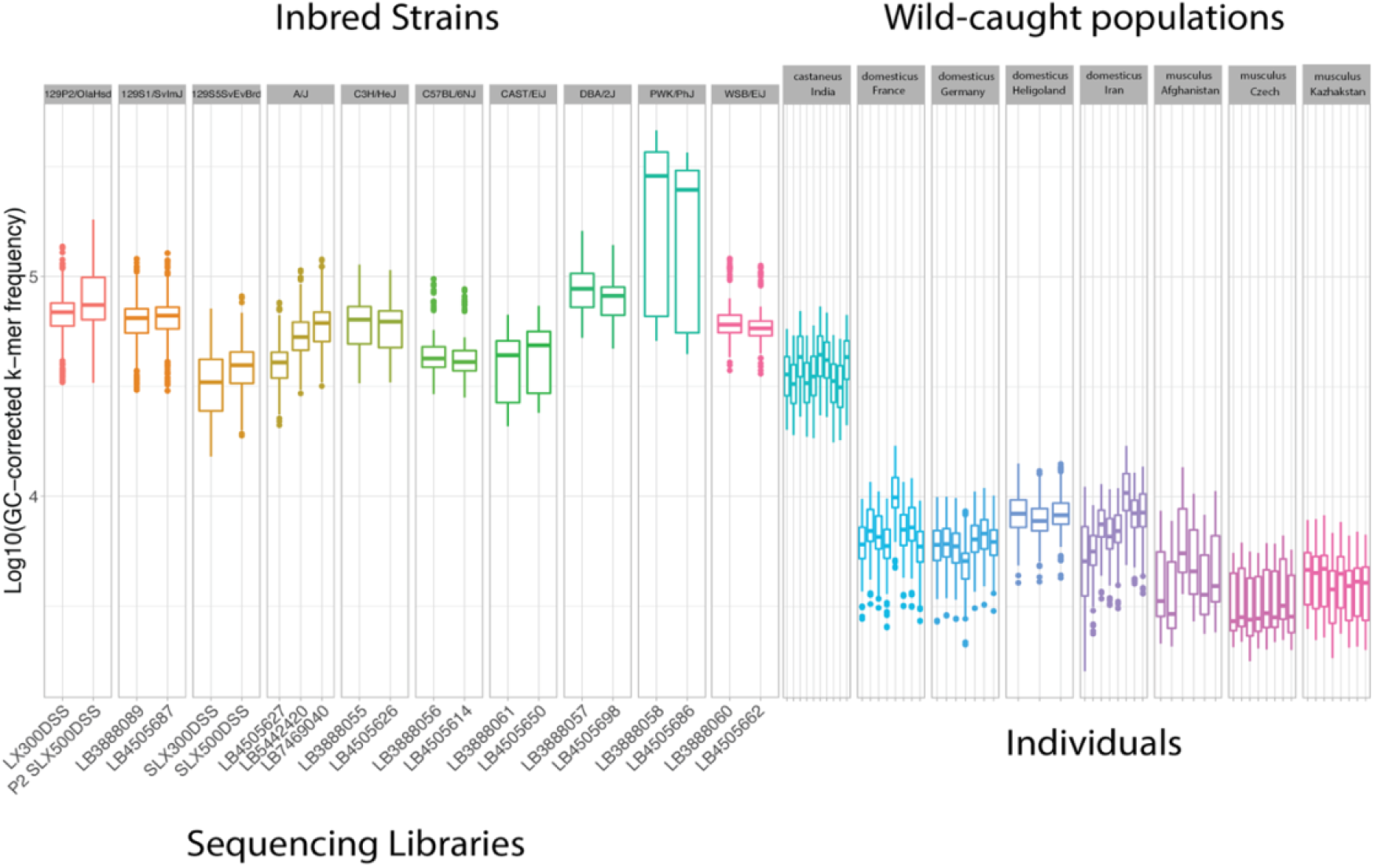
Centromere consensus 31-mer frequencies exhibit low variance between independent sequencing libraries and among wild-caught individuals from a population. Boxplots reveal the distribution of minor centromere satellite 31-mer frequencies for individual sequencing libraries or wild-caught individuals.

### Quantifying within genome centromere satellite diversity

To quantify centromere satellite diversity within a genome, we computed the average number of sequence differences between independent satellite repeats, a metric we term the *centromere diversity index* (CDI). We first mapped sequenced reads to the major and minor centromere consensus sequences using bwa version 0.7.9 (Li & Durbin, 2010). We then partitioned reads using samtools version 1.8 (Li et al., 2009) based on (1) whether they mapped to the major or minor satellite, (2) whether they mapped to the forward or reverse strand to prevent comparing sequences to their reverse complement, and (3) their mapped position along the consensus sequence. For each pair of reads mapping to an identical site in the same orientation on the major or minor satellite sequence, we computed the average number of observed sequence differences, *d_ij_*. We then derived the CDI by averaging over all *N* tested read pairs:

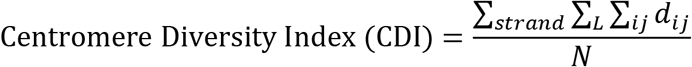

where L is the length of the satellite repeat unit (*L*=120 and *L*=234 for the minor and major satellites respectively).

### Quantifying consensus centromere satellite polymorphisms

To summarize the sequence polymorphism landscape across centromere satellite repeats, we identified *k*-mers with a fixed edit distance (h) of the minor and major satellite consensus sequence. For *k*=15, we allowed *h* ≤ 2, and for *k*=31 we allowed *h* ≤ 5. We used the frequencies of these relaxed edit-distance *k*-mers, in conjunction with their positions across their respective satellite consensus sequences, to derive a vector of relative nucleotide probabilities for each position in the satellite consensus sequence. At a given position, we computed the total frequency of *k*-mers with an “A”, “C”, “G”, or “T” at the focal position. These per-nucleotide *k*-mer frequencies were then converted to relative probabilities summing to one and used to populate a 4×*N* “polymorphism matrix” for each analyzed sample, where *N*=120 for the minor satellite sequence and *N*=234 for the major satellite. Note that this approach ignores the contribution of indel mutations to sequence polymorphism at centromere satellite repeats. We then compared the percentage of non-consensus nucleotides for each strain across the minor and major consensus satellite sequence.

### Phylogenetic analysis of centromere diversity

A total of 56,500,187 high quality SNPs from 12 inbred *Mus musculus* genomes were used to construct a Maximum Likelihood (ML) tree was constructed using RAxML version 8.2.12 (Stamatakis, 2014). The inbred strain SPRET/EiJ was included as an outgroup. An initial set of 20 ML trees was constructed using the GTRGAMMA substitution model. These trees were used as input for subsequent branch length and topology refinements in order to estimate the tree with the highest likelihood. We then used *GTRCAT* to derive bootstrap support values for the best ML tree, with the number of random seeds set to 12345.

We applied Lynch’s phylogenetic comparative method to estimate the phylogenetic heritability of centromere satellite copy number and CDI (Lynch, 1991). Under a neutral (*i.e*., Brownian motion) model of evolution, the extent of phenotypic divergence between species should be proportional to their genetic divergence. We computed the phylogenetic variance-covariance matrix from the *Mus musculus* ML phylogeny and then used this matrix to estimate the proportion of variation in both major and minor satellite copy number and CDI that is explained by the underlying tree. This quantity was then divided by the total variance in satellite copy number and CDI to calculate the phylogenetic heritability 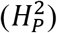 of each diversity parameter. The significance of observed 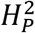 values was assessed by an *ad hoc* permutation test. We shuffled observed satellite copy number and CDI values across the tree tips and then re-estimated 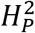 on each permuted dataset. Empirical *P*-values were determined from the quantile position of the observed 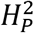 value along the distribution of 1000 permuted 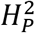 values.

All analyses were performed in R using the Analysis of Phylogenetics and Evolution (ape v5.3) package (Paradis & Schliep, 2019).

### Animal husbandry and ethical commitment

All animal experiments were approved by the Jackson Laboratory’s Animal Care and Use Committee and carried out in compliance with National Institutes of Health guidelines.

The following inbred mouse strains were obtained from The Jackson Laboratory: CAST/EiJ, LEWES/EiJ, PWK/PhJ, WSB/EiJ, and PAHARI/EiJ. Mice were housed in a low barrier room and provided food and water *ad libitum*. Mice were euthanized by CO_2_ asphyxiation or cervical dislocation in accordance with recommendations from the American Veterinary Medical Association.

### Mouse embryonic fibroblasts cultures

Primary mouse embryonic fibroblasts (MEFs) were isolated from E12.5-E13.5 embryos of male and female mice from four inbred strains: CAST/EiJ, LEWES/EiJ, PWK/PhJ, and WSB/EiJ. MEFs were cultured in MEF media composed of Dulbecco’s Modified Eagle medium (DMEM) supplemented with 10% FBS (Lonza), 100ug/mL Primocin (Invivogen) and 1xGlutaMAX (Thermo Fisher Scientific /GIBCO). MEFs were cultured in 150mm tissue culture-treated plates (Thermo Fisher Scientific) at 37°C in a humidified atmosphere with 5% CO_2_.

### Metaphase chromosome spreads and FISH

MEFs were used for the preparation of metaphase spreads. Briefly, MEFs were cultured to ~ 80% confluency at 37°C in a humidified atmosphere with 5% CO_2_ in MEF media. Cells were subsequently serum starved on MEF media without FBS and exposed to 0.02 ug/ml Colcemid (Thermo Fisher Scientific/GIBCO) for 12 hours to synchronize and arrest cells in metaphase. MEFs were subsequently shaken off and resuspended in hypotonic solution (56 mM KCl) for 60 min. The harvested cells were then gradually fixed in 3:1 Methanol:Glacial Acetic Acid under constant agitation. Cells were pelleted by centrifugation, the fixative decanted off, and re-fixed for a total of 3-4 times. Following the final fixation round, cells were suspended in a 1-2 mL volume of fixative and dropped onto slides from a height of ~1m. Slides were allowed to air dry for approximately 10 minutes and then stored at −20C until hybridization.

Commercially synthesized oligos corresponding to the *Mus musculus* major satellite, *Mus musculus* minor satellite, and the predicted *Mus pahari* centromere sequences were fluorescently labelled via nick translation (Supplementary Table 2). Briefly, 250 - 1000 ng of synthesized DNA was combined with nick translation buffer (200 mM Tris pH 7.5, 500 mM MgCl_2_, 5mM Dithiothreitol, 500 mg/mL Bovine Serum Albumin), 0.2 mM dNTPs, 0.2mM fluorescent nucleotides, 1U DNAse (Promega) and 1U DNA Pol I (Thermo Fisher Scientific). Three fluorescent nucleotides were used: Fluorescein-12-dUTP (Thermo Fisher Scientific), ChromaTide Texas Red-12-dUTP (Thermo Fisher Scientific/Invitrogen), and Alexa Fluor 647-aha-dUTP (Thermo Fisher Scientific/Invitrogen). The reaction mixture was incubated at 14.5 C for 90 minutes, and then terminated by addition of 10mM EDTA. Probes ranged from 50-200 bp in size, as assessed by gel electrophoresis.

**Supplementary Table 2: Sequences and primers used for FISH experiments.**

Probes were used in FISH reactions on MEF metaphase cell spreads. Probes were denatured in hybridization buffer (50% formamide, 10% Dextran Sulfate, 2x saline-sodium citrate (SSC), mouse Cot-1 DNA) at 72°C for 10 min and then allowed to re-anneal at 37°C until slides were ready for hybridization. Slides were dehydrated in a sequential ethanol series (70%, 90%, 100%; each 5 min) and dried at 42°C. Slides were then denatured in 70% formamide/2x SSC at 72°C for 3 min, and immediately quenched in ice cold 70% ethanol for 5 minutes. Slides were subjected to a second ethanol dehydration series (90%, 100%; each 5 min) and air dried. The probe hybridization solution was then applied to the denatured slide. The hybridized region was then cover-slipped and sealed with rubber cement. Hybridization reactions were allowed to proceed overnight in a humidified chamber at 37°C. After gently removing the rubber cement and soaking off coverslips, slides were washed 2 times in 50% formamide/2x SSC followed by an additional 2 washes in 2x SSC for 5 min at room temperature. Slides were counterstained in 0.05ng/mL DAPI (Thermo Fisher Scientific/Invitrogen) for 10 min and air dried at room temperature. Lastly, slides were mounted with ProLong Gold AntiFade (Thermo Fisher Scientific/Invitrogen) and stored at −20C until imaging.

### Image capture and fluorescence intensity quantification

FISH reactions were imaged at 63x magnification on a Leica DM6B upright fluorescent microscope equipped with fluorescent filters and a cooled monochrome 2.8 megapixel digital camera. Images were collected at a plane with maximal intensity using consistent exposure settings across slides (DAPI at 20ms, TxRed at 50ms and FITC at 200ms). The mean intensities of FISH signals at each centromere were calculated in areas drawn around centromeres based on thresholding with background subtraction. Signals were quantified from all centromeres within a cell (n = 40). FISH fluorescent intensity signals were collected from two independent cell lines (biological replicates) from each strain and two independent experiments were conducted for each cell line with fluorophores swapped for each sequence (technical replicates). We collected images from 8-10 cells per replicate, amounting to >320 individual centromere measurements per replicate (40 centromeres signal/cell x 8 cells ≥ 320). Differences in fluorescent intensity between strains were assessed by ANOVA (baseR). Fluorescent intensity is represented in arbitrary units (AU).

### Evaluating signals of meiotic drive in the Diversity Outbred mapping population

We utilized genotype probability data from five Diversity Outbred (DO) mapping studies conducted on mouse cohorts from outbreeding generations 6 to 22 (Bult MegaMUGA, Svenson-183 MegaMUGA, Churchill-181 MegaMUGA, Attie-232 GigaMUGA, and Chesler-192 MegaMUGA; all data from https://www.jax.org/research-and-faculty/genetic-diversity-initiative/tools-data/diversity-outbred-database). All DO mice were genotyped at a common set of loci(Churchill, Gatti, Munger, & Svenson, 2012). For mice in each outbreeding generation, we first determined the frequency of each parental haplotype at every genotyped marker. We then looked for linked clusters of markers that exhibit a consistent departure from the expected haplotype frequency (0.125) and that displayed a monotonic increase in the frequency of one or more haplotypes over successive outbreeding generations. Such regions may harbor loci subject to meiotic drive loci.

### Mouse Phenotype Data

Spearman correlation tests were used to test for relationships between chromosome instability phenotypes and estimated centromere satellite copy number across inbred lab strains. Chromosome instability phenotypes were obtained from the Mills1 dataset deposited in the Mouse Phenome Database(Bogue et al., 2020).

## Results

### *k*-mer analysis reveals striking differences in the abundance of centromere satellite repeats across *Mus*

Standard approaches for surveying sequence diversity are not readily extendable to the centromere due to its repeat-rich architecture and gapped status on the current mouse reference assembly. To circumvent these challenges, we employed a *k*-mer based approach to quantify the diversity of satellite DNA in mouse genomes. Our *k*-mer strategy is predicated on the insight that the relative frequency of a given nucleotide word, or *k*-mer, in a shot-gun sequencing library is proportional to its frequency in the parent sample genome. Thus, the observed frequency of a particular *k*-mer within a pool of sequenced reads can be used as a proxy for its relative abundance in a genome. We normalized *k*-mer counts to adjust for potential GC-biases introduced during library preparation, and confirmed through rigorous comparisons of replicate libraries for individual samples that our corrected *k*-mer counts provided a reliable readout of the relative frequency of nucleotide motifs in diverse mouse genomes (Supplementary Figure 1; See Methods).

We first identified the most abundant 15-mers across a subset of 54 diverse mouse genomes. These genomes included common inbred mouse strains, wild-caught mice from multiple populations from each of the three principle house mouse subspecies (*M. m. domesticus*, *M. m. castaneus*, and *M. m. musculus*), and three divergent *Mus* taxa (*M. spretus, M. caroli, and M. pahari*). Consistent with prior reports (Komissarov, Gavrilova, Demin, Ishov, & Podgornaya, 2011), *Mus musculus* minor and major centromere satellite 15-mers were among the most abundant sequences in mouse genomes (top 0.01% of all 15-mers). Interestingly, centromere 15-mers were also among the most differentially abundant 15-mers across diverse *Mus musculus* genomes (Figure 1), hinting at extensive centromere satellite copy number variation.

**Figure 1:**
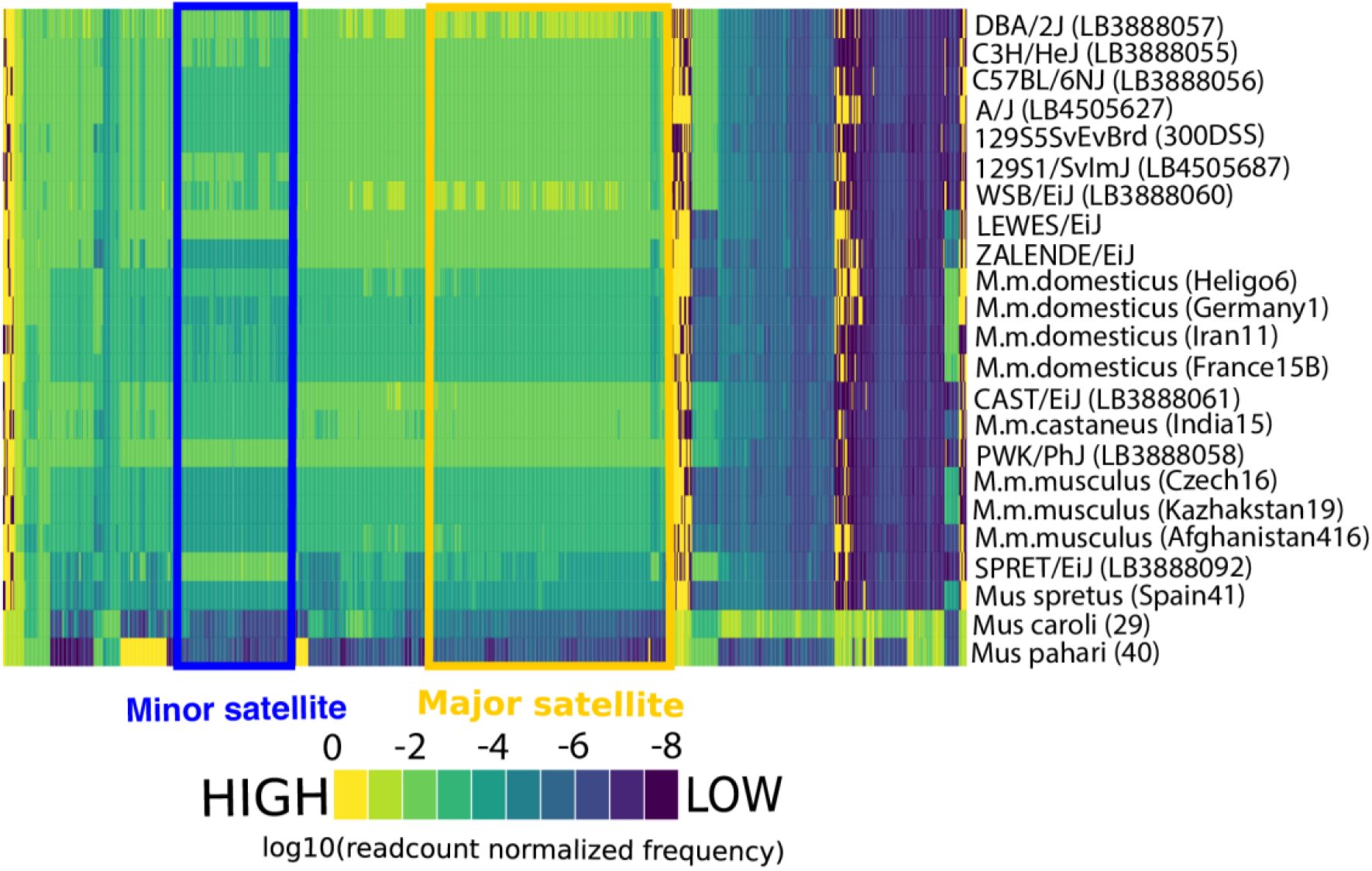
Consensus centromere 15-mers were the most abundant and variable 15-mers in diverse *Mus* genomes. Heatmap displaying the observed frequencies of the 1000 most variable 15-mers (columns) across a sample of diverse *Mus* genomes (rows). Supplementary Figure 4 profiles a larger set of samples (n = 54). The color scale represents the frequency of all centromere satellite 15-mers, normalized by the number of sequenced reads. 15-mers present in the *Mus musculus* minor and major satellite consensus sequences are noted by the blue and yellow boxes, respectively.

*Mus spretus* shares an identical minor satellite consensus sequence with *Mus musculus* (Narayanswami et al., 1992), and exhibited a high abundance of minor satellite centromere 15-mers (Figure 1). In contrast, *Mus caroli* harbors divergent centromere satellite sequences from those in *M. musculus* (Kipling et al., 1995). Expectedly, we found very weak enrichment for *M. musculus* major and minor consensus centromere 15-mer sequences in the*M. caroli* genome. Similarly, we found no enrichment for *M. musculus* major and minor centromere 15-mers in *M. pahari*, suggesting that *M. pahari* centromeres are also defined by a unique and divergent satellite.

**Supplementary Figure 4:**
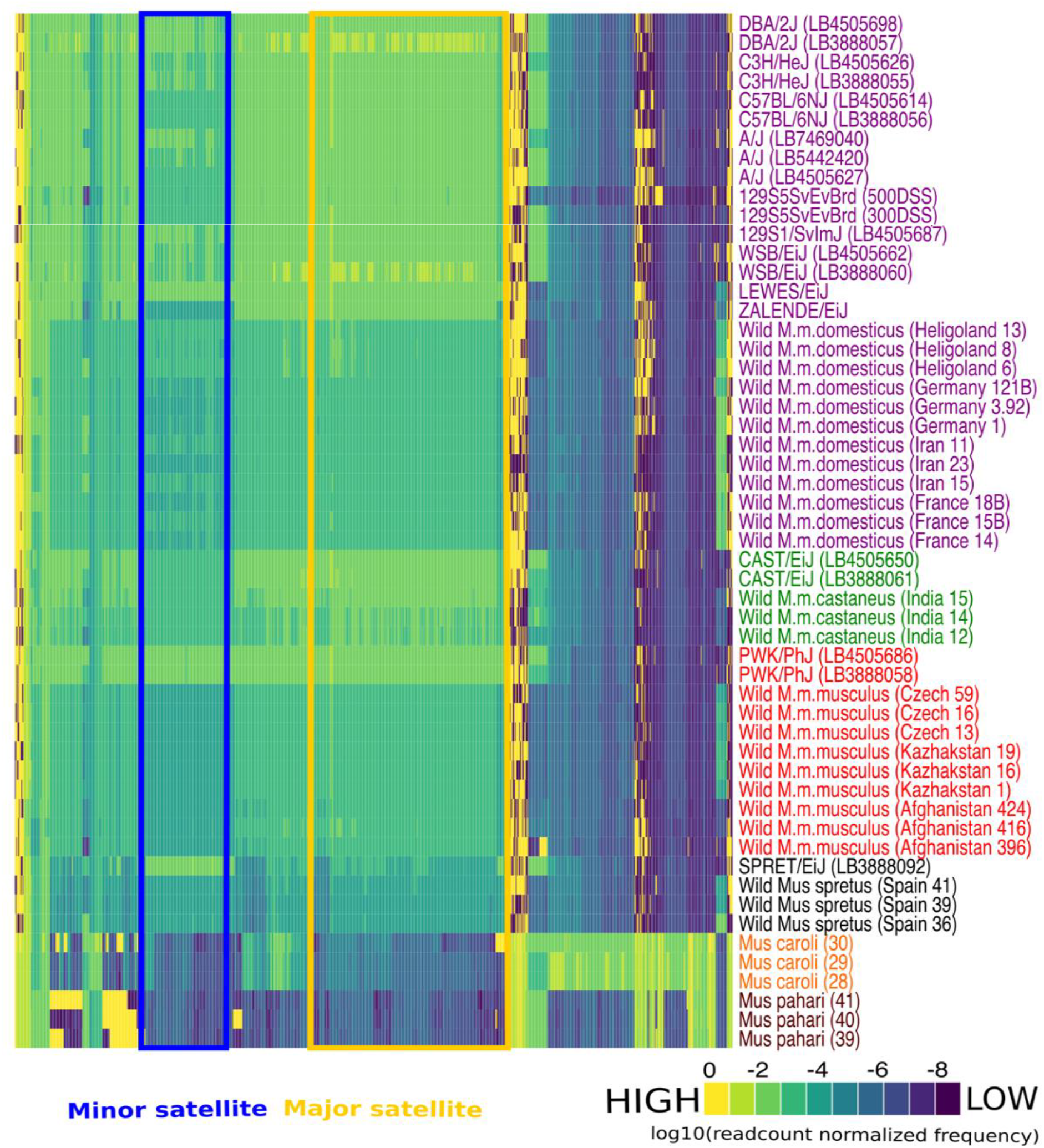
Consensus centromere 15-mers were the most abundant and variable 15-mers in diverse *Mus* genomes. Heatmap displaying the observed frequencies of the 1000 most variable 15-mers (columns) across 54 diverse samples (rows). The color scale represents the frequency of all centromere satellite 15-mers, normalized by the number of sequenced reads. 15-mers present in the *Mus musculus* minor and major satellite consensus sequences are noted by the blue and yellow boxes, respectively.

### Strain and population-level variation in the abundance of the *Mus musculus* consensus centromere satellites

Owing to the high prevalence and striking variability in the abundance of centromere satellite 15-mers among *Mus* genomes, we sought to further define strain, subspecies, and species variation in both major and minor satellite copy number. We compared the GC-corrected frequencies of minor and major satellite 31-mers across *Mus musculus* genomes (Figure 2A) and uncovered several noteworthy trends. For the following observations we present data for 31-mers but observed qualitatively identical trends for 15-mers (Supplementary Figure 5). First, we uncovered a greater abundance of major satellite 31-mers compared to minor satellite 31-mers (Figure 2B; Kruskall-Wallis one-way ANOVA, P < 2.2×10^-16^). This difference is consistent with the known size differences between the major and minor satellite array in *Mus musculus* (Cazaux et al., 2013; Komissarov et al., 2011; Wong & Rattner, 1988). Second, there was greater strain-to-strain variation in the abundance of minor satellite *k*-mers as compared to the major satellite *k*-mers across the inbred strains (Kruskall-Wallis one-way ANOVA d.f. = 13, minor satellite F value = 907.8, major satellite F value = 309.3, P < 2.2×10^-16^). We converted our normalized *k*-mer counts into absolute satellite copy number estimates to approximate strain and subspecies differences in consensus centromere size (see Methods). We estimate between 3100 – 125,000 minor satellite copies and 19,000 – 630,000 major satellite copies in the genomes of 14 inbred *Mus musculus* strains. These estimates include only exact matches to the consensus satellite sequences and ignore the potential presence of other sequence elements that modify centromere size differences between samples. Nonetheless, the 40-(33-) fold range in absolute minor (major) consensus satellite copy number suggests remarkable differences in centromere size between closely related inbred *Mus musculus* strains. Third, inbred strains harbored higher satellite 31-mer frequencies than wild-caught mice (Figure 2; Student’s *t*-test = 212.76; P < 2.2×10^-16^). Indeed, PCA analyses of minor and major 31-mer frequencies across diverse *Mus musculus* samples identified inbred versus wild (i.e., outbred) as the major axes of differentiation (Supplementary Figure 6). This outcome is not an artifact of systematically undercounting centromeric *k*-mers with sequence mismatches to the consensus, as we also observe a reduced fraction of reads mapping to the centromeric consensus in wild mice compared to inbred strains (Supplementary Figure 2). We speculate that inbreeding may lead to the expansion of centromeric repeats in house mice, similar to observations and an earlier proposal for maize (Schneider, Xie, Wolfgruber, & Presting, 2016).

**Figure 2:**
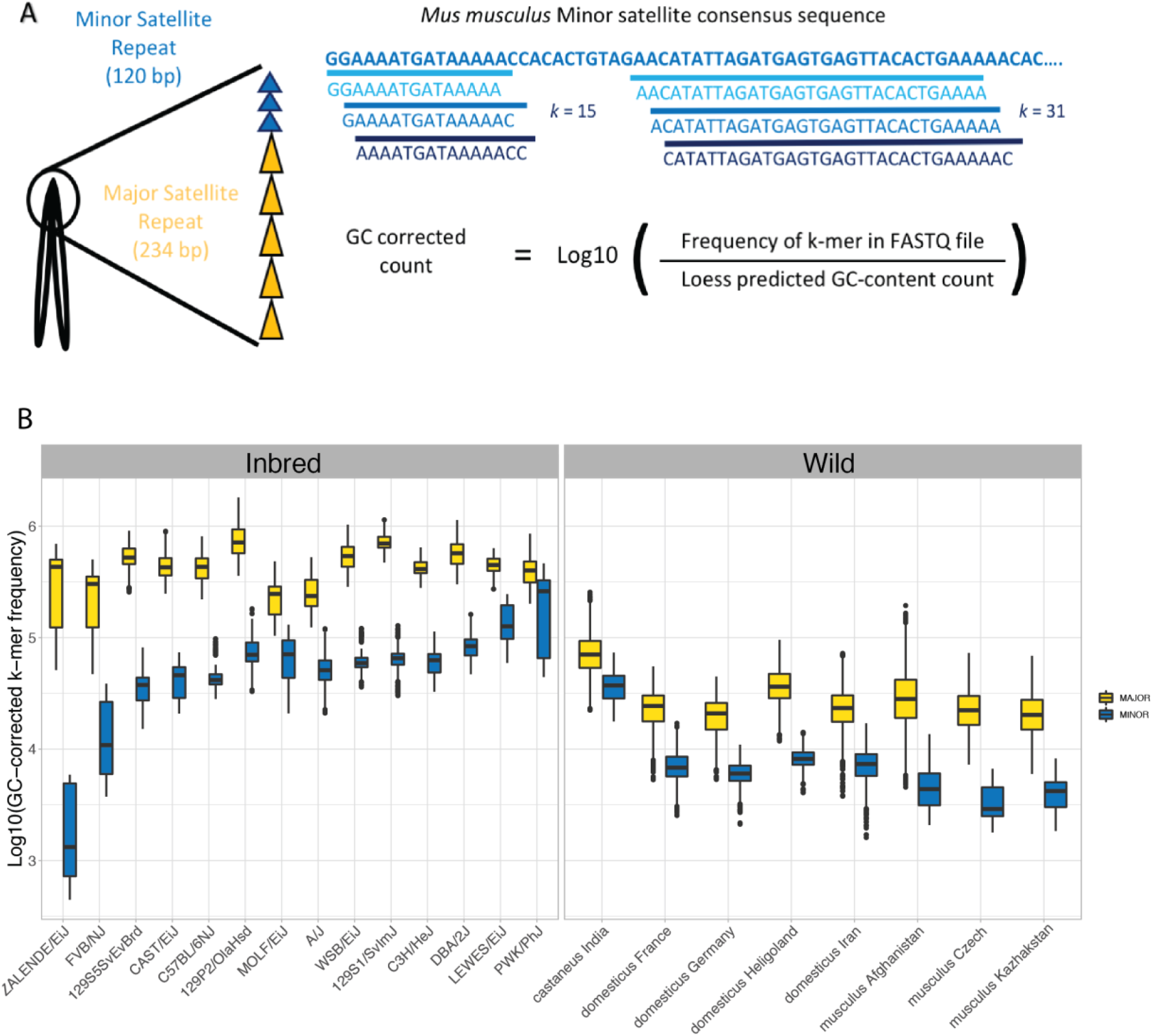
Significant differences in consensus centromere satellite copy number across *Mus musculus*. (A) Schematic overview of the approach used to quantify the frequencies of *k*-mers in centromeric satellite repeats. (B) Boxplots showing the distribution of major (yellow) and minor (blue) satellite 31-mer frequencies across inbred strains and wild-caught mouse populations.

**Supplementary Figure 5:**
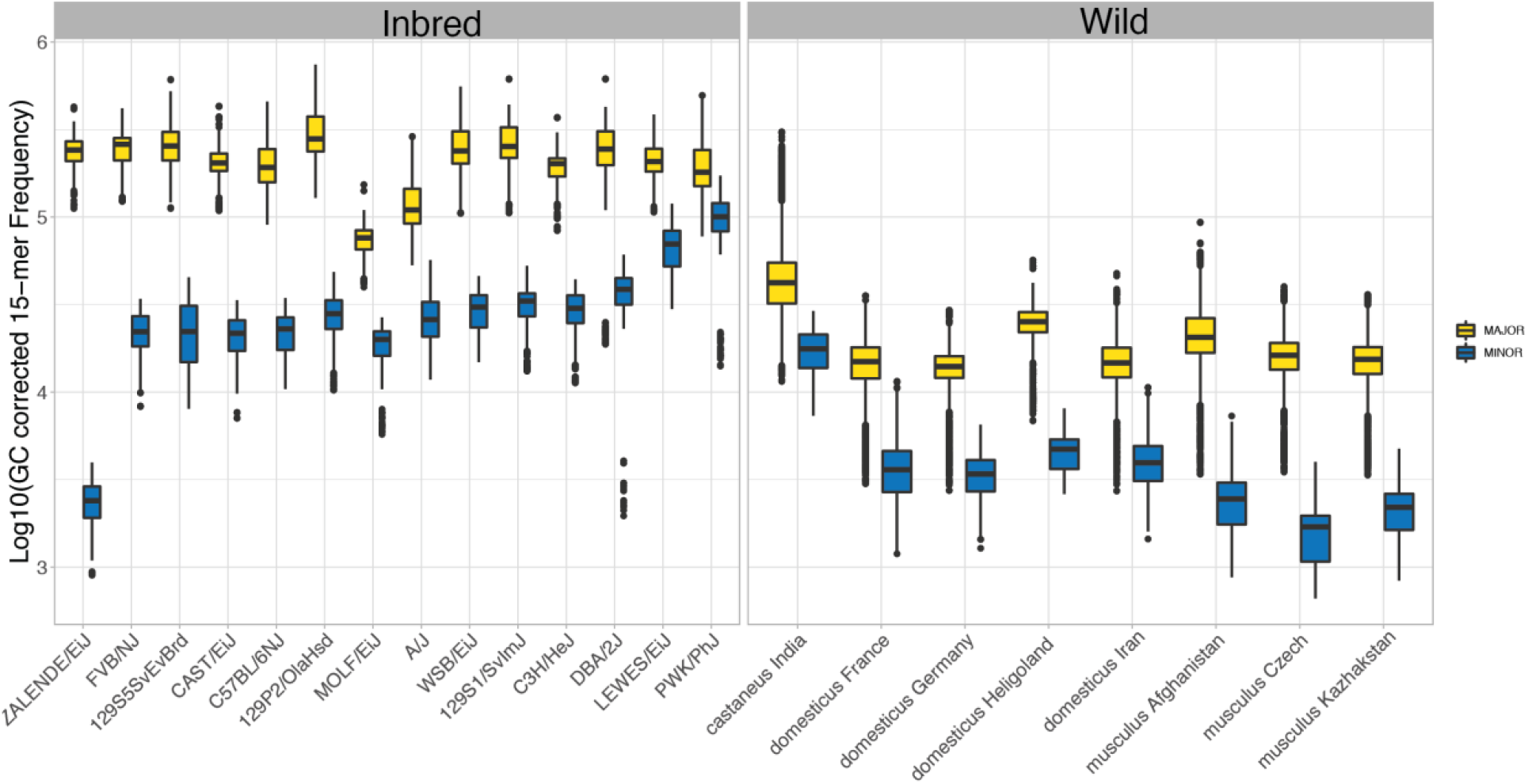
Variation in centromere consensus 15-mer frequencies across diverse *Mus musculus* genomes. Boxplots reveal the distribution of major (yellow) and minor (blue) satellite 15-mer frequencies across inbred strains and wild-caught mouse populations.

**Supplementary Figure 6:**
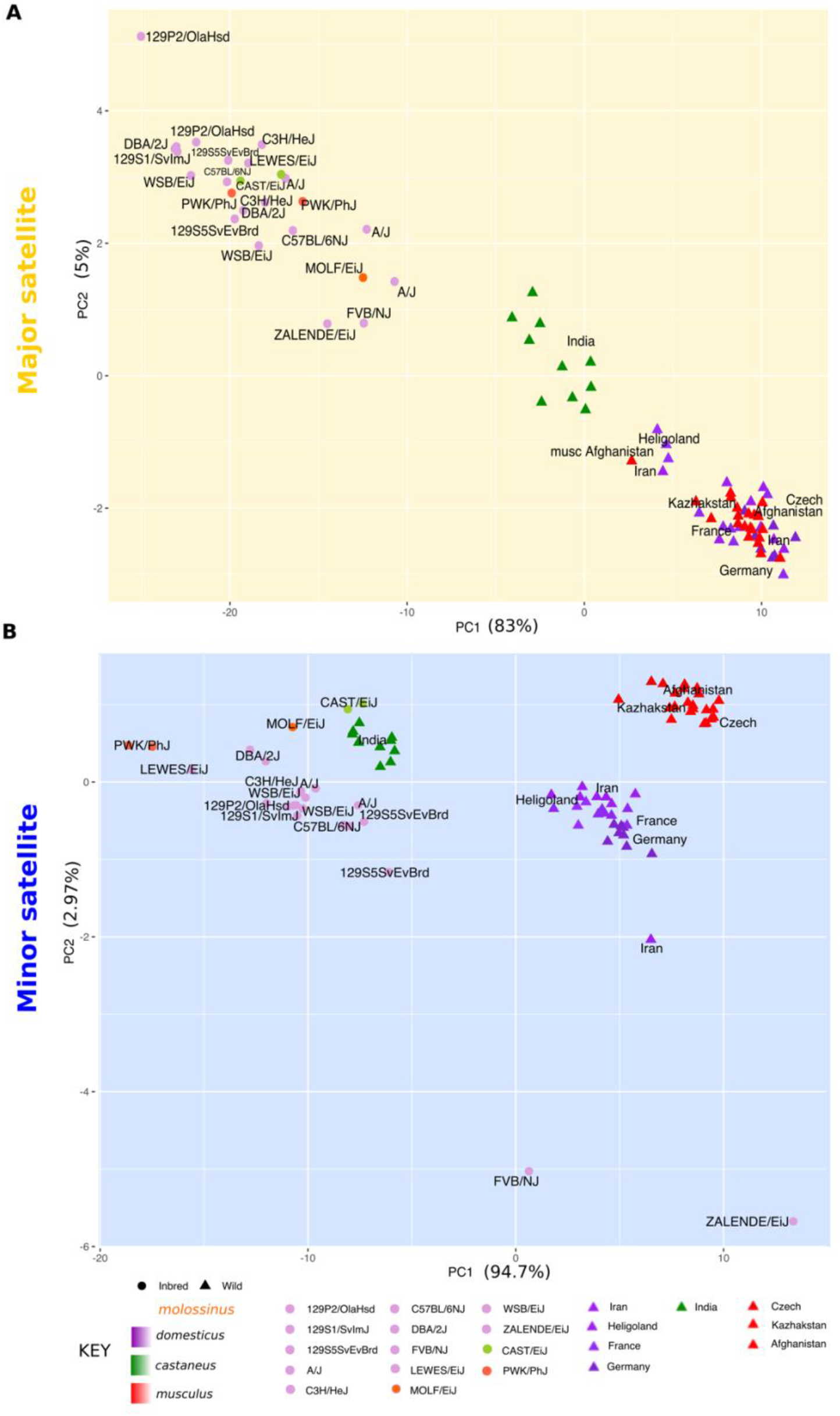
Inbred strains and wild-caught mice exhibit distinct centromere *k*-mer frequencies. Principal component analysis of (A) major and (B) minor satellite centromere 31-mer frequencies in inbred strains and wild-caught *M. musculus* samples.

### Cytogenetic validation of strain differences in consensus centromere satellite abundance

We used quantitative FISH (qFISH) to validate our *k*-mer based estimates of strain variation in consensus centromere satellite abundance. We focused on a subset of strains that encompass a range of estimated minor satellite copy numbers and span three principle house mouse subspecies: CAST/EiJ (*M. m. castaneus*), WSB/EiJ (*M. m. domesticus*), LEWES/EiJ (*M. m. domesticus*), and PWK/PhJ (*M. m. musculus*) (Figure 2B). We observed strong overall concordance between relative copy number and qFISH signals at both the minor and major centromere satellites (Figure 3B). Notably, both methods yielded a similar rank order of strains with respect to the minor satellite abundance (median fluorescent intensity ranking in arbitrary units WSB/EiJ = 3440 < CAST/EiJ = 3780 < LEWES/EiJ = 3820 < PWK/PhJ = 7055). Interestingly, in WSB/EiJ and LEWES/EiJ, several chromosomes consistently showed a higher minor satellite signal intensity relative to other chromosomes (Figure 3A). This observation contrasted more uniform intensity of the minor satellite signal across all chromosomes in CAST/EiJ and PWK/PhJ (Figure 3A). These findings point to chromosome-specific minor satellite accumulation and/or loss in some *Mus musculus*, highlighting an additional dimension of centromere diversity.

**Figure 3:**
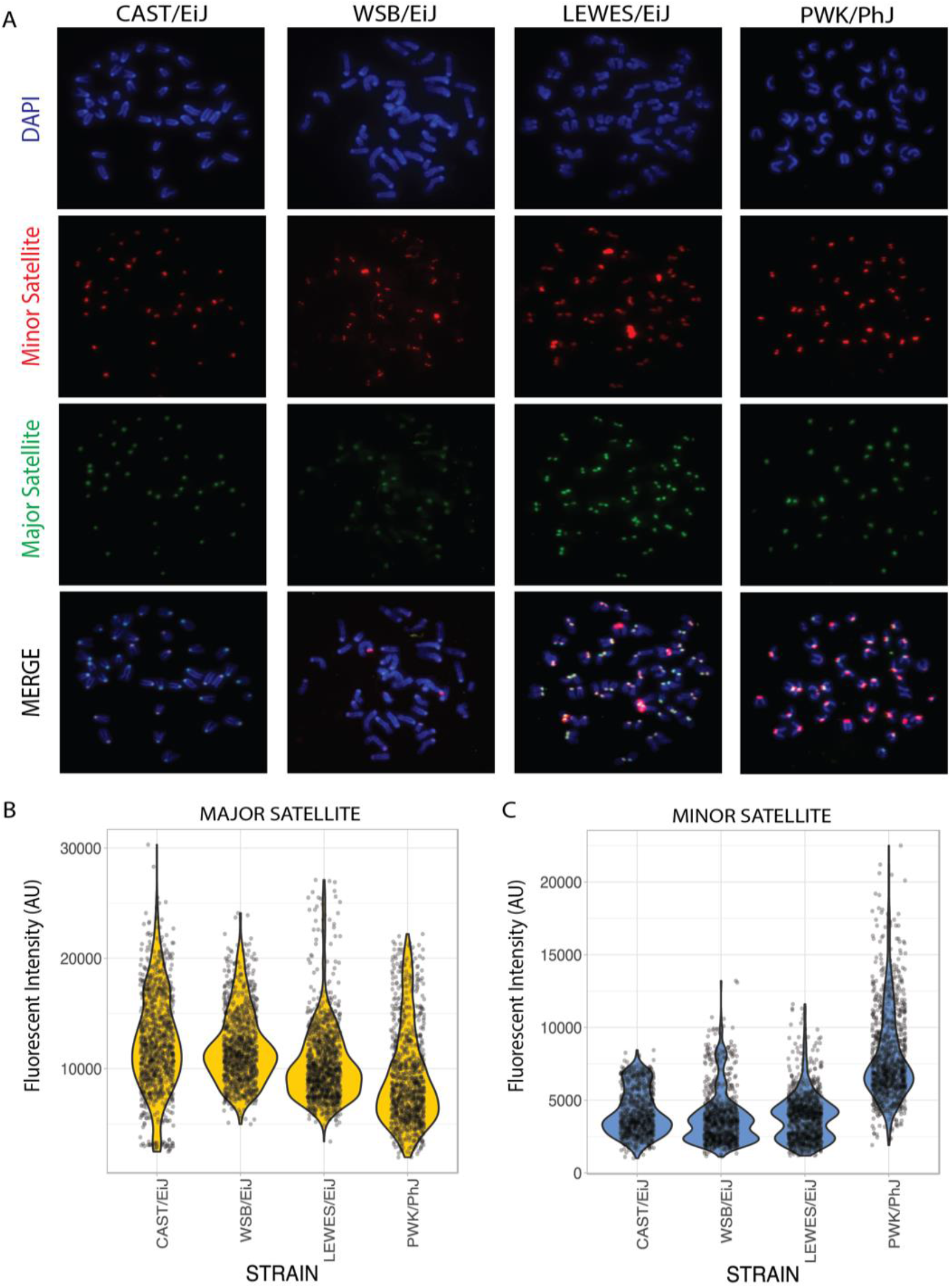
Quantitative FISH reveals consensus centromere satellite copy number variation across inbred mouse strains. (A) Representative FISH images for four genetically diverse inbred strains: CAST/EiJ, WSB/EiJ, LEWES/EiJ, and PWK/PhJ (B and C) Quantification of fluorescent intensity using DNA probes derived from the (B) major and (C) minor centromeric satellite repeats across inbred strains. Points correspond to fluorescent intensity measurements for a single chromosome. A minimum of 40 centromeres from 36 cells were examined per strain. Fluorescent intensity is represented in arbitrary units (AU).

### Centromere satellite repeat heterogeneity at the strain and population level

Using *k*-mers with exact matches to the consensus sequences, we have uncovered significant variation in consensus centromere satellite copy number across diverse *Mus musculus* samples. However, focusing on *k*-mers with exact matches to the consensus limits our ability to discover and analyze centromere repeat diversity within a genome. To study this important class of centromere variation, we calculated the average number of pairwise sequence differences between centromere satellite repeats in each *Mus musculus* sample (see Methods). We refer to this metric as the *centromere diversity index* (CDI). The minor satellite CDI across inbred *Mus musculus* strains is lower than in wild-caught *Mus musculus* mice (inbred range: 12.1-23.0, wild range: 24.8-35.6), revealing greater homogeneity of minor satellite repeats in inbred strains compared to wild-caught mice. For both inbred and wild mice, the average minor satellite CDI (inbred = 17.6, wild = 30.7) is slightly higher than the major satellite CDI (inbred = 16.9, wild = 28.7), despite the increased length and greater genomic abundance of the major satellite. Based on these findings, we conclude that the mouse minor satellite harbors appreciably higher sequence diversity than the major satellite.

We next assessed the relationship between centromere diversity and consensus satellite copy number. There is an overall negative correlation between satellite copy number and CDI for both the minor and major satellite repeats (Figure 4A and B; minor satellite: Spearman’s rho=-0.40, *P*= 0.08; major satellite: Spearman’s rho = −0.7, *P*= 0.001). Samples with high satellite copy number had more homogenous repeats, whereas samples with lower copy numbers had higher repeat heterogeneity. This relationship is largely driven by the striking distinction between wild-caught mice and inbred strains. Relative to inbred strains, wild-caught mice harbored smaller and more diverse centromere arrays. The similarity in minor satellite size and diversity in inbred strain CAST/EiJ and wild-caught *M. m. castaneus* represents one possible exception to this pattern (Figure 4A). Together, these trends suggest that phenomena specific to inbred strain genomes (or, potentially, the very process of inbreeding itself) may have influenced centromere architecture in pronounced ways.

**Figure 4:**
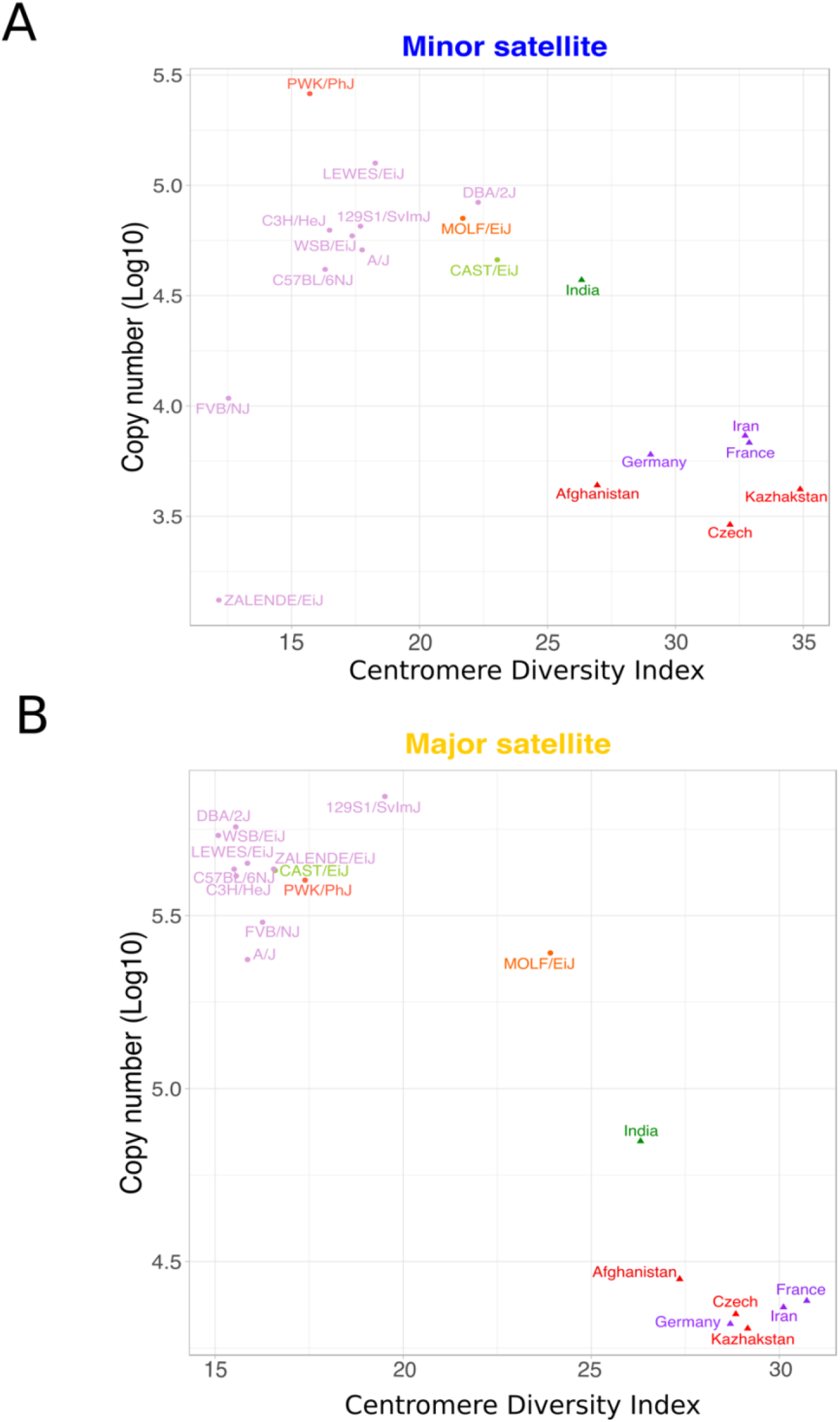
Negative correlation between centromere satellite copy number and sequence diversity in *Mus musculus*. Estimated centromere satellite copy number and centromere diversity index for the (A) minor or (B) major satellite sequence. Copy number was estimated from the median frequency of consensus centromere 31-mers in each sample. The three primary house mouse subspecies are denoted by different colors: red - *M. m. musculus*, purple - *M. m. domesticus*, and green - *M. m. castaneus*, orange – *M. m. molossinus*. Shapes distinguish inbred strains (circles) from wild-caught mice (triangles).

### Sequence landscape of *Mus musculus* satellite diversity

Our CDI measure captures overall centromere satellite diversity within single genomes, but does not pinpoint specific satellite sequence positions that subject to high variability. To investigate the landscape of sequence polymorphisms along the major and minor centromere satellite repeats, we relaxed the criterion for perfect *k*-mer matching by considering all 15- and 31-mers with ≤2 and ≤5 mismatches, respectively, from the *Mus musculus* centromere satellite consensus sequences. These relaxed edit-distance *k*-mers can be unambiguously assigned to positions in the minor and major satellite consensus sequences, allowing us to quantify the proportion of *k*-mers harboring nucleotide mismatches at each position. Using the percentage of non-consensus nucleotides at each position, we then identified sites with variable nucleotide usage across samples.

Overall, sequence diversity is not uniformly distributed across the minor and major satellite sequences, but instead restricted to a limited number of sites that are variable between genomes (Figure 5). Despite its smaller size, the minor satellite harbors more sites with at least 20% non-consensus nucleotide usage than the major satellite (107 versus 79; Figures 5A and 5B). Although divergence from the satellite consensus is concentrated at a minority of sites, different samples vary in the frequency of non-consensus nucleotides present at a given position. For example, LEWES/EiJ, WSB/EiJ and 129S1/SvImJ have similar centromere diversity indices (CDI=17−18), but their minor satellite sequence landscapes are distinct from each other (Figure 5A).

**Figure 5:**
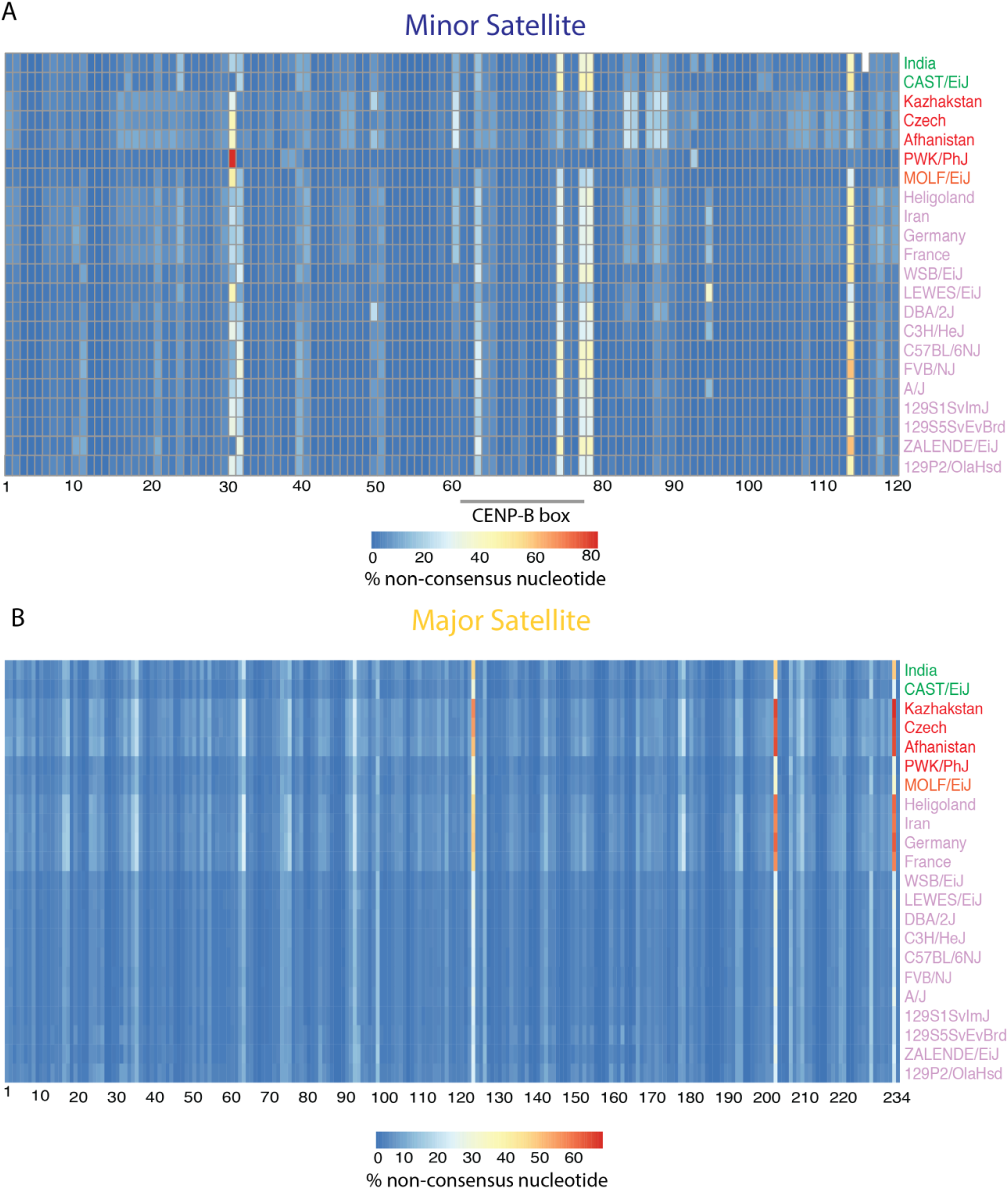
Landscape of nucleotide variation across centromeric satellite repeats. Heatmap of non-consensus nucleotide usage for positions in the (A) minor satellite consensus sequence and (B) major satellite consensus sequence. Each row corresponds to a single sample with sample names color-coded by subspecies origin: green – *M. m. castaneus*, red – *M. m. musculus*, purple – *M. m. domesticus*, orange – *M. m. molossinus*.

Intriguingly, three positions within the CENP-B binding motif of the minor satellite show high levels of nucleotide variability among *Mus musculus*. CENP-B binding is important but dispensable for kinetochore assembly and chromosome segregation (Hudson et al., 1998). Whether observed variants in the CENP-B box lead to differences in the binding efficiency of CENP-B across *Mus musculus* remains unknown. Similarly, the potential functional significance of nucleotide variation at other satellite positions will require future investigation.

We also uncover clear differences in the satellite sequence landscape between wild-caught mice and inbred strains. On average, inbred strains have lower rates of non-consensus nucleotide usage (minor satellite 2.9-4.9%; major satellite 3.6-4.4%) compared to wild-caught mice (minor satellite 4.8-6.8%; major satellite 5.7-6.5%). This finding aligns with the higher CDI observed in wild-caught compared to inbred mice, suggesting that wild-caught *M. musculus* have more diverse and heterogenous centromere satellites than inbred strains.

### Rapid evolution of minor satellite copy number and repeat heterogeneity

To investigate how centromere architecture evolves in house mice, we analyzed the distribution of centromere diversity metrics in a phylogenetic framework (Lynch, 1991). Using phylogenetic comparative methods, we quantified the proportion of strain-to-strain variation in major and minor satellite copy number and CDI that is explained by the underlying strain phylogeny (Figure 6). We limited this analysis to inbred strains owing to the large contrast in centromere architecture between inbred and wild-caught mice. Inclusion of both types of animals would likely mask any legitimate phylogenetic signals within either group. We further excluded the inbred strain ZALENDE/EiJ from this analysis, as it harbors multiple Robertsonian chromosomal rearrangements associated with the loss of centromere satellite sequence that could similarly confound the detection of phylogenetic patterns.

**Figure 6:**
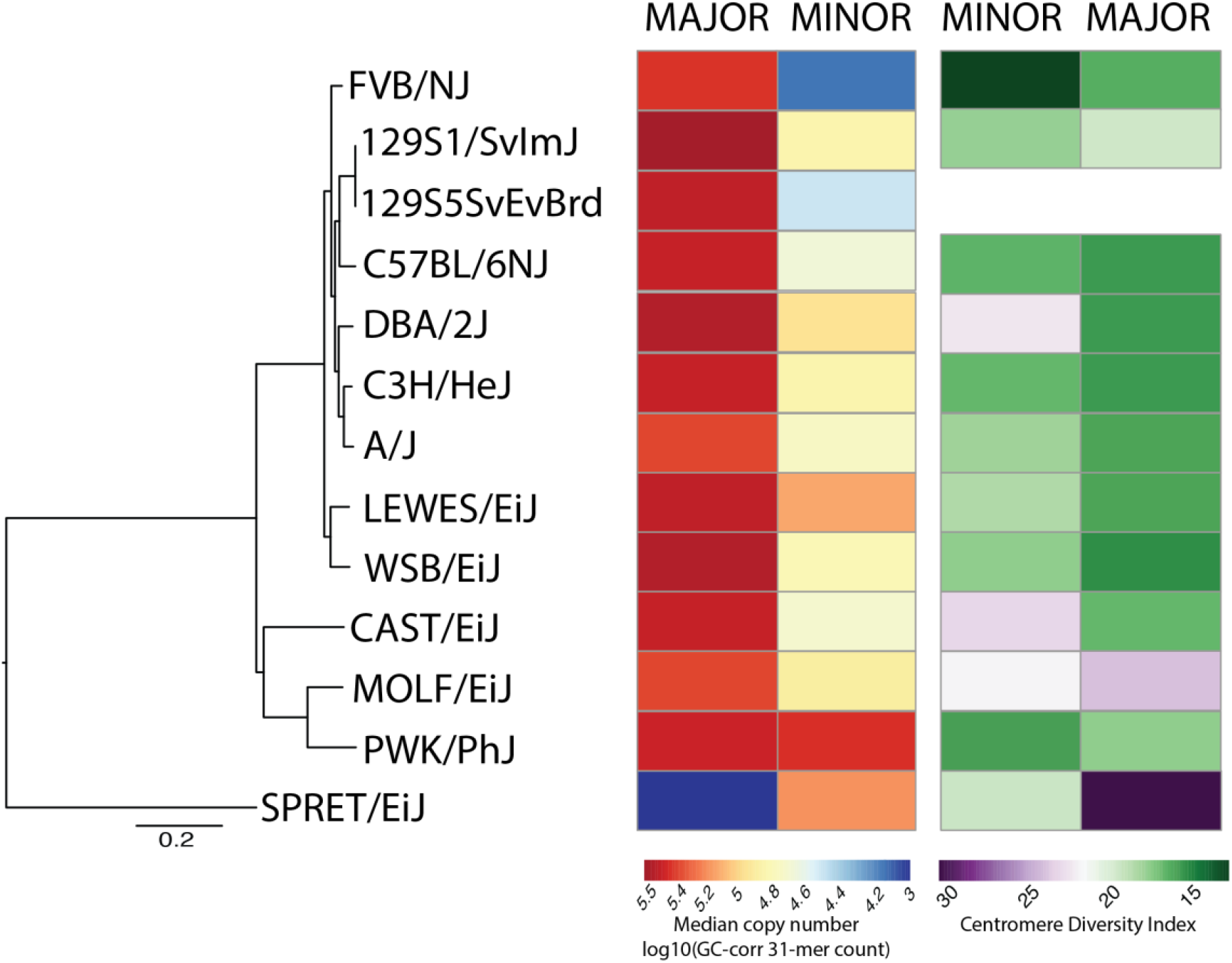
Phylogenetic distribution of centromere satellite copy number and satellite diversity in inbred mice. Maximum likelihood phylogenetic tree for 12 inbred house mouse strains and the outgroup, SPRET/EiJ. For each strain and for both the major and minor satellites, estimated satellite copy number (median 31-mer frequency) and satellite heterogeneity (CDI) are indicated by boxes shaded according to the corresponding color scales.

If variation in satellite copy number or satellite heterogeneity (*i.e*. CDI) is well-predicted from the evolutionary relationships among inbred strains and *M. musculus* subspecies, these metrics should exhibit a high phylogenetic heritability, 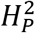. In contrast, if centromere satellite copy number or levels of satellite heterogeneity evolve at exceptionally high rates, these measures of centromere variation should exhibit a weak phylogenetic signal (*i.e*., low 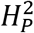). Consistent with this latter prediction, the phylogenetic heritability of both minor satellite copy number 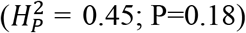 and CDI 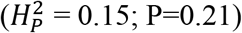 was low, and not significantly different from zero. Evidently, both measures of minor satellite variation evolve sufficiently rapidly to outpace signals of strain relatedness. In contrast, variation in both major satellite copy number 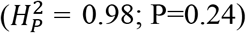 and CDI 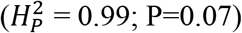 exhibited a high, albeit non-significant, phylogenetic heritability. Although modest sample sizes limit the power of this analysis, the absolute differences in the 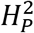 estimates between the minor and major satellites suggests that these two centromere satellites are evolving via distinct regimes, potentially mediated by differences in selective pressures or mutational mechanisms.

### Assessing the phenotypic consequences of centromere diversity in *Mus musculus*

Centromere integrity is essential for genome stability and if not maintained can lead to cancer and infertility (Aldrup-MacDonald et al., 2016; Barra & Fachinetti, 2018; Hudson et al., 1998; Régnier et al., 2005; Zhang et al., 2016). We next asked whether observed centromere satellite diversity influences the stability of genome transmission. Using publicly available phenotype data from the Mouse Phenome Database (https://phenome.jax.org/) we searched for correlations between centromere satellite copy number and micronuclei formation, a hallmark of chromosome instability (Lee et al., 2013; Luzhna, Kathiria, & Kovalchuk, 2013). We found no significant correlation between this measure of genome stability and either major or minor satellite consensus copy number (Supplementary Figure 7). These results suggest no strong functional link between centromere size and this measure of chromosome instability. However, small sample sizes, uncertainty in our copy number estimates, and imprecision in the chromosomal instability phenotype may conceal true biological associations.

**Supplementary Figure 7:**
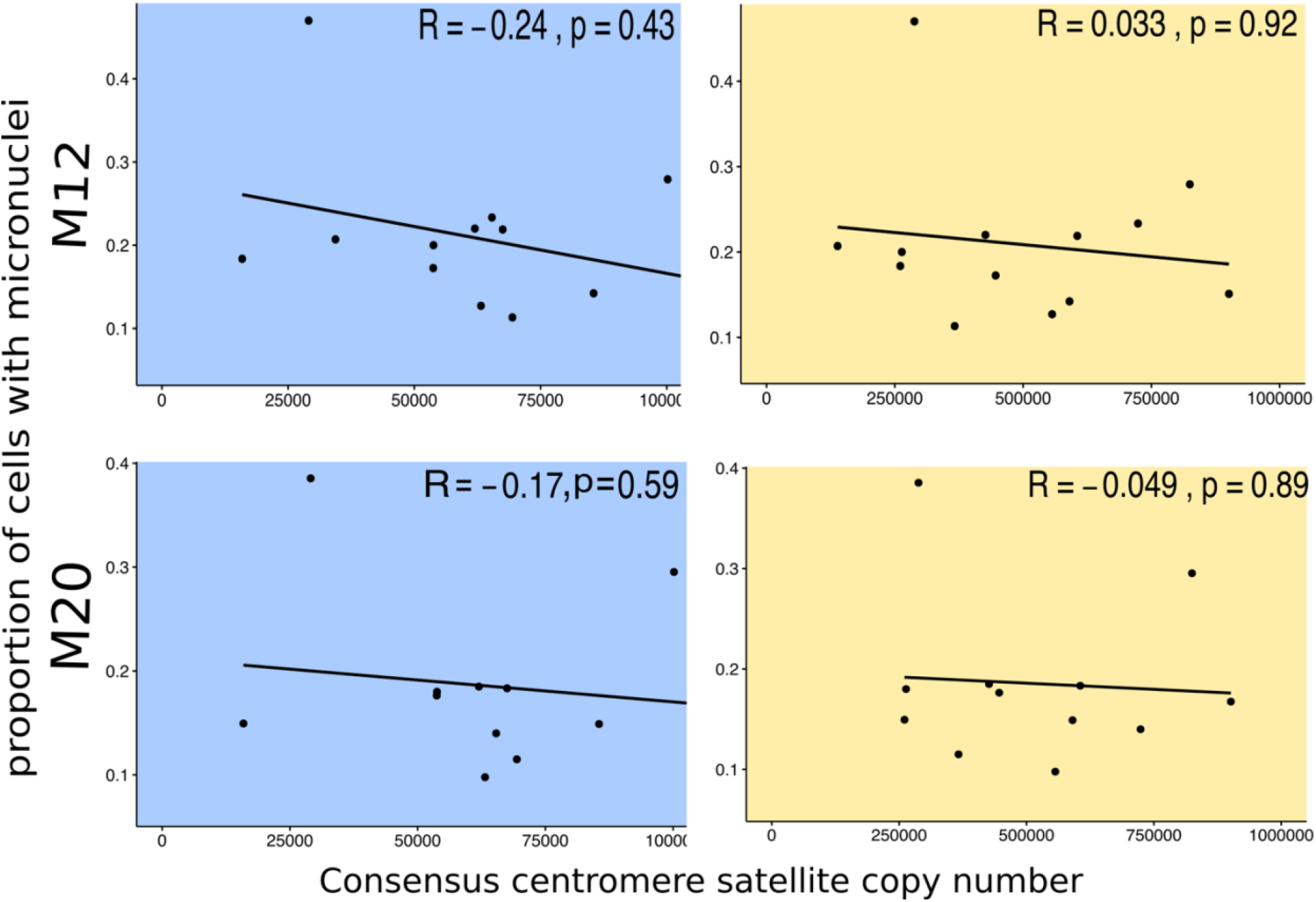
No correlation between micronuclei frequency and centromere satellite consensus copy number. Spearman correlations between the proportion of peripheral blood cells (red blood cells and micronuclei) with micronuclei and median minor (left) or major (right) satellite 31-mer frequencies. The proportion of cells with micronuclei was determined for 12-month-old mice (top) and 20-month-old mice (bottom).

Centromeres are reservoirs for the accumulation of selfish drive elements (Chmátal et al., 2014; Kursel & Malik, 2018) that can hijack the inherent asymmetry of female meiosis to bias their own transmission into the oocyte (Kursel & Malik, 2018; Malik, 2009). We next asked whether centromere satellite copy number differences among inbred strains lead to systematic meiotic drive in diverse mouse populations. We profiled datasets from the Diversity Outbred (DO) mouse population (Churchill et al., 2012), a heterogenous stock population founded from 8 strains with distinct centromere satellite copy number states. We scanned genotypes of DO mice from 15 successive generations for evidence of the over-transmission of centromere-proximal alleles from one (or more) founder strain(s). We found no evidence for non-Mendelian transmission of centromere-proximal regions in the DO (Supplementary Figure 8). This result suggests (i) the absence of centromere-mediated meiotic drive in this complex population, (ii) the lack of power to detect weak drive signals, (iii) that drive is influenced by multiple genetic factors (Didion et al., 2016), or (iv) that aspects of centromere architecture other than minor satellite copy number may be critical for defining drive potential.

**Supplementary Figure 8:**
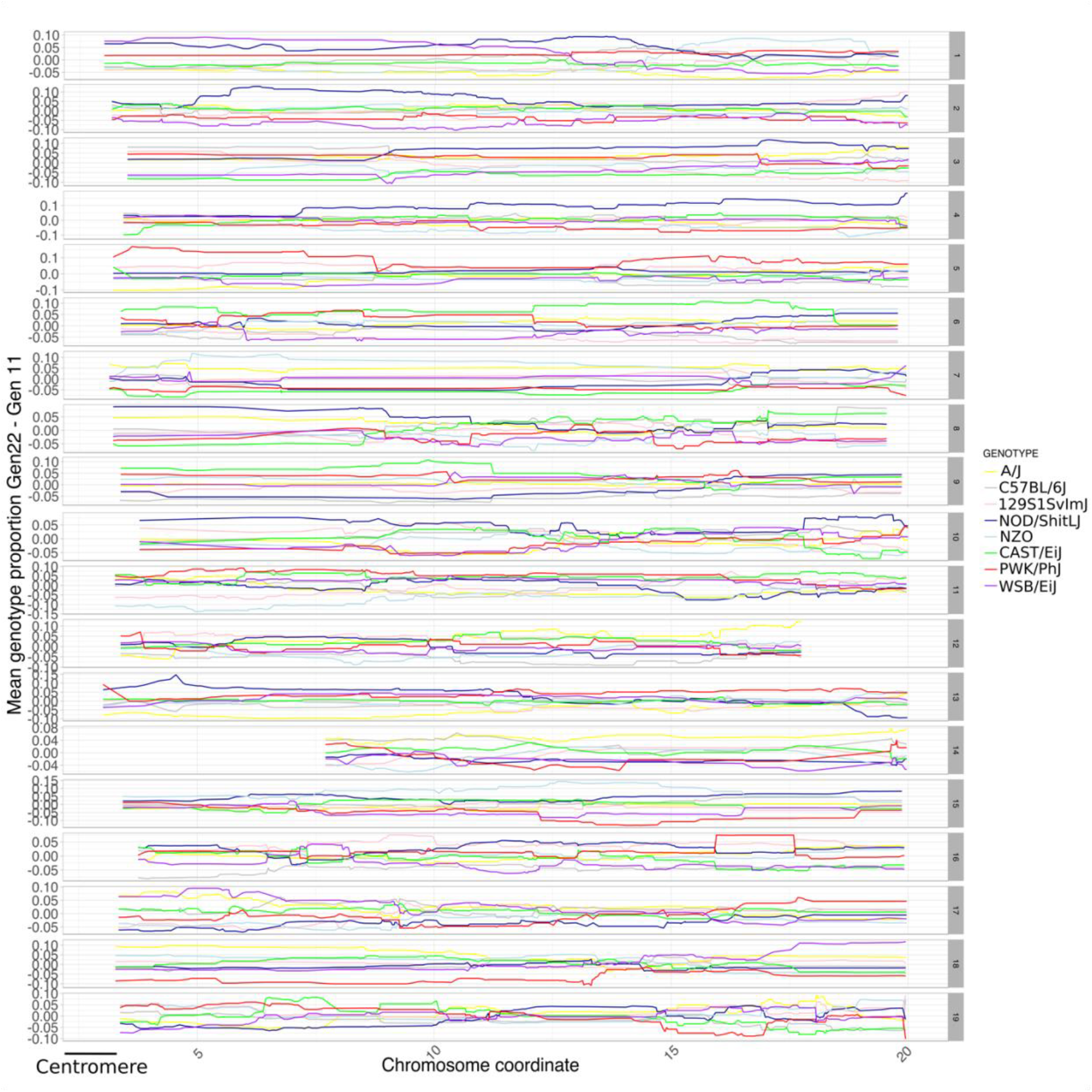
Haplotype frequencies at centromere-proximal regions in the Diversity Outbred populations are not consistent with strong centromere drive. Chromosome coordinates of genotyped markers in megabases (Mb) are provided on the x-axis. The difference in the frequency of each strain haplotype between generation 22 and generation 11 is shown on the y-axis. Line colors correspond to each of the 8 DO founder strains.

## Discussion

Evolutionary theory predicts that genomic regions with key cellular roles should exhibit reduced rates of evolution in order to preserve their biological function. Centromeres are paramount for chromosome segregation and the maintenance of genome stability, but, paradoxically, centromere satellite sequences are known to evolve rapidly between species (Alkan et al., 2011; Feliciello, Akrap, Brajković, Zlatar, & Ugarković, 2014; Garrido-Ramos, 2017; Smith, 1976). Despite this knowledge, comparatively little is known about the extent of centromere variation over shorter evolutionary timescales, including at the population level. Here, we developed a powerful *k*-mer based workflow for quantifying centromere satellite copy number and sequence diversity from whole genome sequence data. We apply this analytical framework to 100 genomes from diverse inbred and outbred mice to characterize multiple dimensions of mouse centromere variation.

Our analyses present several notable advances. First, whereas prior studies have used genomic methods to survey the diversity of centromere satellite sequences across divergent taxa (Melters et al., 2013), our study represents one of the first efforts to uncover the short-term evolutionary dynamics of telocentric centromeres and the first in-depth study of centromere diversity in house mice. Second, whereas earlier studies of centromere diversity in mice applied qualitative approaches to small cohorts of inbred strains (Aker & Huang, 1996), our study provides quantitative estimates of both centromere copy number and diversity across inbred strains and wild-caught mice. Third, by situating our findings in a phylogenetic framework, we show that centromere satellite copy number and heterogeneity are volatile and rapidly evolving properties of centromere architecture. Lastly, by pursuing these investigations in a well-established model system with rich phenotyping resources, our work presents an initial functional exploration of observed centromere diversity and encourages further investigations of its functional effects on the fidelity and dynamics of chromosome segregation.

We discovered key differences in the mode and rate of evolution of the *Mus musculus* major and minor satellite sequences. Minor satellite arrays exhibited more extreme variation in copy number and CDI in comparison to the major satellite arrays (Figure 2B and Figure 4). Using phylogenetic comparative methods, we further showed that the rate of evolution in minor satellite copy number and CDI is sufficiently rapid to erode signals of strain relatedness. In contrast, major satellite copy number variation exhibits a stronger phylogenetic signal, although we lack sufficient power to obtain statistically significant results (Figure 6). These differences between the *Mus musculus* major and minor satellite repeat are presumably due to their distinct biological functions. The major satellite repeat forms the pericentromeric heterochromatin and is responsible for the establishment and maintenance of sister chromatid cohesion (McKinley & Cheeseman, 2016). The minor satellite repeat binds to CENP-A, a specialized centromeric histone variant responsible for kinetochore complex specification and assembly (McKinley & Cheeseman, 2016). In many animal species, CENP-A is rapidly evolving, which imposes a complementary selection pressure on the centromere satellite sequence to ensure protein-DNA compatibility (Henikoff, Ahmad, & Malik, 2001; Malik & Henikoff, 2001; Talbert, Bryson, & Henikoff, 2004). The CENP-A amino acid sequence is perfectly conserved among *M. musculus* subspecies, but sequence diversity at the centromere satellite could influence the efficiency of CENP-A binding, with potential downstream consequences for kinetochore assembly and chromosome segregation (Iwata-Otsubo et al., 2017; Sullivan, Chew, & Sullivan, 2017). In contrast, the *M. musculus* major satellite sequence does not serve as a sequence substrate for kinetochore proteins. The co-evolutionary dynamics between the minor satellite DNA and CENP-A have likely contributed to the accelerated evolution of the minor satellite relative to the major satellite.

Our work also identified surprising differences in centromere satellite architecture between wild-caught and inbred mice. Wild-caught mice exhibit lower major and minor centromere satellite copy numbers and greater satellite heterogeneity than the inbred strains (Figure 5). Similar observations have been previously reported for centromeres in inbred and outbred maize (Schneider et al., 2016). Together, these findings suggest that the inbreeding process itself might drive the homogenization of satellite arrays and facilitates the fixation of larger centromeres. Indeed, prior studies have established that larger centromeres may recruit more kinetochore proteins than smaller centromeres, enabling larger centromeres to selfishly bias their own segregation into the oocyte during asymmetric female meiosis, a process known as centromere drive (Akera, Trimm, & Lampson, 2019; Chmátal et al., 2014; Iwata-Otsubo et al., 2017). In the context of inbreeding, such “strong centromeres” should be rapidly fixed. Recurrent bouts of de novo centromere expansion and fixation could lead to rapid, run-away amplification of centromere satellites in inbred strains compared to wild-caught animals. Thus, centromere size and repeat heterogeneity within inbred strains may not faithfully capture the native state of *M. musculus* centromeres. Future investigations that chronical changes in centromere size from the earliest stages of inbreeding onward could provide a real-time window into the mutational processes that promote this architectural shift.

Our analyses define the extent of centromere copy number and sequence diversity in diverse inbred strains, begging investigation into the phenotypic consequences of this variation. As an initial attempt to address this outstanding challenge, we looked for correlations between satellite copy number and a phenotype proxy for chromosome instability: the frequency of spontaneous micronuclei formation in peripheral blood cells. We observed no significant relationship between these variables, although the tested phenotype – spontaneous micronuclei formation – is likely an imprecise measure of centromere-mediated genome instability (Luzhna et al., 2013). Furthermore, our analysis was limited to a small number of inbred strains with available published data, and is underpowered to find small to moderate strength genotype-phenotype correlations. We also tested whether variation in minor satellite copy number leads to centromere drive in an outbred mouse population from eight inbred strains with variable minor satellite copy number (Churchill et al., 2012). We found no evidence for strong non-Mendelian transmission of centromere-proximal variants although again, our analysis likely suffers from a lack of statistical power to find weak to moderate drive signals. By providing the first quantitative estimates of centromere satellite diversity in a panel of widely used inbred strains, our investigation critically informs strain choice for future studies that aim to rigorously and explicitly test how centromere diversity influences the fidelity of chromosome segregation and genome stability.

Ultra-long read sequencing technologies are now enabling sequence-level resolution of mammalian centromeres (Jain et al., 2018; Longsdon et al., 2020; Miga, 2019; Miga et al., 2020). However, the high cost of these methods and their labor-intensive analyses put their use out of reach for most investigators and effectively limit the number of population samples that can be analyzed. Our powerful *k*-mer-based workflow for assaying the architectural and sequence diversity of centromeres circumvents these critical limitations and is readily extendable to large numbers of genomes. Using only short read data in public repositories, our work has provided key evolutionary insights into the scope of population and subspecies variation across house mouse centromeres, establishing the needed foundation for functional tests of centromere diversity in an important biomedical model system.

## Acknowledgements

We thank Drs. Mary Ann Handel (JAX), Laura Reinholdt (JAX), and Christopher Baker (JAX) for their comments while preparing the manuscript.

